# G-PROTEIN COUPLED RECEPTOR 19 (GPR19) KNOCKOUT MICE DISPLAY SEX-DEPENDENT METABOLIC DYSFUNCTION

**DOI:** 10.1101/2022.10.29.514377

**Authors:** Bellina A.S. Mushala, Bingxian Xie, Ian J. Sipula, Michael W. Stoner, Dharendra Thapa, Janet R. Manning, Paramesha Bugga, Amber M. Vandevender, Michael J. Jurczak, Iain Scott

**Author notes:** To whom correspondence should be addressed: Iain Scott, PhD, FAHA, FCVS, Department of Medicine, University of Pittsburgh, BST E1259, 200 Lothrop Street Pittsburgh, PA, 15261. Division of Exercise Physiology, School of Medicine, West Virginia University, Morgantown, WV, 26506.

## Abstract

G-protein coupled receptors (GPCRs) mediate signal transduction from the cellular surface to intracellular metabolic pathways. While the function of many GPCRs has been delineated previously, a significant number require further characterization to elucidate their cellular function. G-protein coupled receptor 19 (GPR19) is a poorly characterized class A GPCR which has been implicated in the regulation of circadian rhythm, tumor metastasis, and mitochondrial homeostasis. In this report, we use a novel knockout (KO) mouse model to examine the role of GPR19 in whole-body metabolic regulation. We show that loss of GPR19 promotes increased energy expenditure and decreased activity in both male and female mice. However, only male GPR19 KO mice display glucose intolerance in response to a high fat diet (HFD). Loss of GPR19 expression in male mice, but not female mice, resulted in diet-induced hepatomegaly, which was associated with decreased expression of key fatty acid oxidation genes in male GPR19 KO livers. Overall, our data suggest that loss of GPR19 impacts whole-body energy metabolism in diet-induced obese mice in a sex-dependent manner.

## 1. INTRODUCTION

As the largest family of membrane proteins, G-protein coupled receptors (GPCRs) transduce extracellular signals to drive intracellular physiological responses. GPCRs mediate signaling involved in vision, olfaction, behavior, neurotransmission, immunity, and metabolic homeostasis, and consequently their dysfunction is implicated in many disease states ^[1] [2] [3] [4]^. There are over 800 GPCRs encoded by the human genome which are categorized into five classes - *Glutamate, Rhodopsin, Adhesion, Frizzled/Taste2*, and *Secretin -* based on their sequence and structural similarity ^[5] [6] [7]^. The class A Rhodopsin-like receptors are the largest, most diverse, and well-studied GPCR group, containing more than 75% of the identified receptors ^[5]^. Despite their ubiquitous expression and physiological significance, more than 140 GPCRs have undetermined endogenous ligands and/or biological functions, and are thus termed “orphan GPCRs” ^[8] [9]^.

G-coupled protein receptor 19 (GPR19) is a class A orphan GPCR that was first identified from a human genome expressed sequence tag (EST) database ^[10]^, displaying abundant expression in the brain observed during mouse embryogenesis ^[11]^. A recent report further demonstrated that GPR19 is expressed in the suprachiasmatic nucleus of the adult hypothalamus, and may be involved in the regulation of circadian rhythms ^[12]^. In addition to the brain, GPR19 is differentially expressed in several peripheral tissues including the heart, liver, kidney, and testis ^[11] [13] [14]^. A few studies have linked elevated GPR19 expression to tumor progression and metastasis, with these downstream signaling cascades affecting mitochondrial homeostasis ^[15] [16] [17]^. However, there is no current evidence of any pathophysiological relevance for GPR19 in systemic or tissue-specific metabolism. In this study, we characterized the metabolic phenotype of both lean and diet-induced obese wildtype and GPR19 knockout (KO) mice by examining whole-body and tissue-specific effects. Our results suggest that GPR19 deficiency impairs whole-body metabolism in a sex-dependent manner, and may serve a central role in the development of obesity-related metabolic dysfunction.

## 2. RESULTS

### 2.1 GPR19 deficiency modifies whole body metabolism

Previous studies have shown that the orphan G-protein coupled receptor (GPCR) GPR19 is expressed in various tissues, with a predominant role in brain physiology ^[10] [11] [12] [18]^. However, the *in vivo* metabolic function of GPR19 has not yet been defined. To assess this, we subjected wildtype (WT) and whole-body GPR19 knockout (KO) mice to metabolic cage analysis under serial low-fat (LFD) and high-fat (HFD) fed conditions. As expected, HFD induced significant increases in total body weight (Fig. 1A,B) in both males and females. However, significant GPR19-dependent increases in fat mass were only observed in female mice (Fig. 1B). The loss of GPR19 expression increased total energy expenditure in male mice under LFD conditions (Fig. 1C), and in female mice under both LFD and HFD conditions (Fig. 1D), suggesting a potential role for GPR19 as a modulator of energy homeostasis.

**Figure 1.**
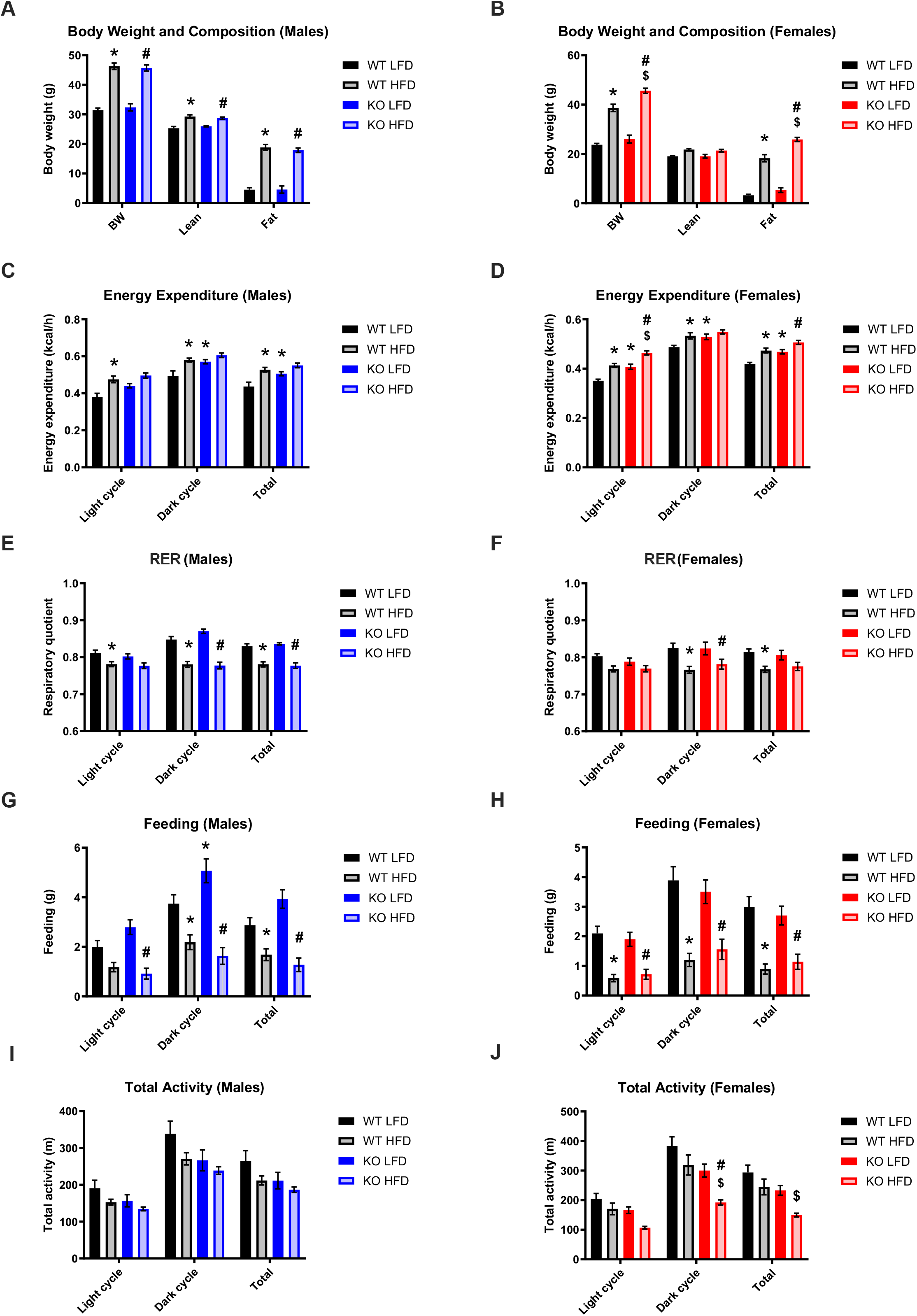
GPR19 deficiency modifies whole body metabolism. Male and female wildtype or GPR19 KO mice were fed a low fat diet (LFD) and subject to metabolic cage analysis. The same mice were then placed on a high fat diet (HFD) for 10 weeks, after which a repeat metabolic cage was performed. (A,B) Body weight and composition, (C,D) energy expenditure, (E,F) respiratory quotient (RQ), (G,H) feeding, and (I,J) total activity were measured in males and females, respectively. * = P < 0.05 vs WT LFD, # = P < 0.05 vs KO LFD, $ = P < 0.05 vs WT HFD, N = 7-9.

The respiratory exchange ratio (RER), derived from the ratio of carbon dioxide produced (VCO_2_) to oxygen consumed (VO_2_), was used to determine changes in whole-body fuel substrate utilization. HFD reduced RER in both male (Fig. 1E) and female (Fig. 1F) mice, suggesting an increased preference for fatty oxidation; however, these changes were not GPR19 dependent. Under HFD conditions, feeding was significantly reduced as expected in male (Fig. 1G) and female mice (Fig. 1H) compared to LFD controls. However, there was a significant increase in feeding activity observed in male KO mice compared to WT mice under LFD conditions during the dark cycle (Fig. 1G). Finally, loss of GPR19 expression in male mice led to a non-significant trend towards decreased activity (Fig. 1I), while in female mice, the loss of GPR19 expression significantly decreased total activity under HFD conditions (Fig. 1J). Overall, loss of GPR19 expression promoted a general increase in energy expenditure, coupled with a general decrease in total activity, suggesting a shift in whole-body energy metabolism.

### 2.2 Glucose homeostasis is impaired in male GPR19 KO mice following exposure to a high fat diet

To determine the effect of GPR19 knockout on whole-body glucose homeostasis, we performed intraperitoneal glucose tolerance tests (IPGTTs) in male and female mice. Under LFD conditions, there were no differences in glucose tolerance in either male (Fig. 2A) or female (Fig. 2D) mice. However, following HFD feeding, loss of GPR19 expression significantly increased circulating glucose levels in male mice (Fig. 2B), but not in female mice (Fig. 2E). As expected, the glucose area under the curve (AUC), an index of whole-body glucose excursion, was significantly increased with high-fat feeding in both WT and KO mice compared to LFD controls (Fig. 2C,F). Male glucose AUC was 31% higher in HFD-fed KO mice compared to WT mice under the same conditions (Fig. 2C), with no significant differences observed in female mice (Fig. 2F). Together, these data suggest that the loss of GPR19 expression attenuates whole-body glucose clearance in diet-induced obesity in a sex-dependent manner.

**Figure 2.**
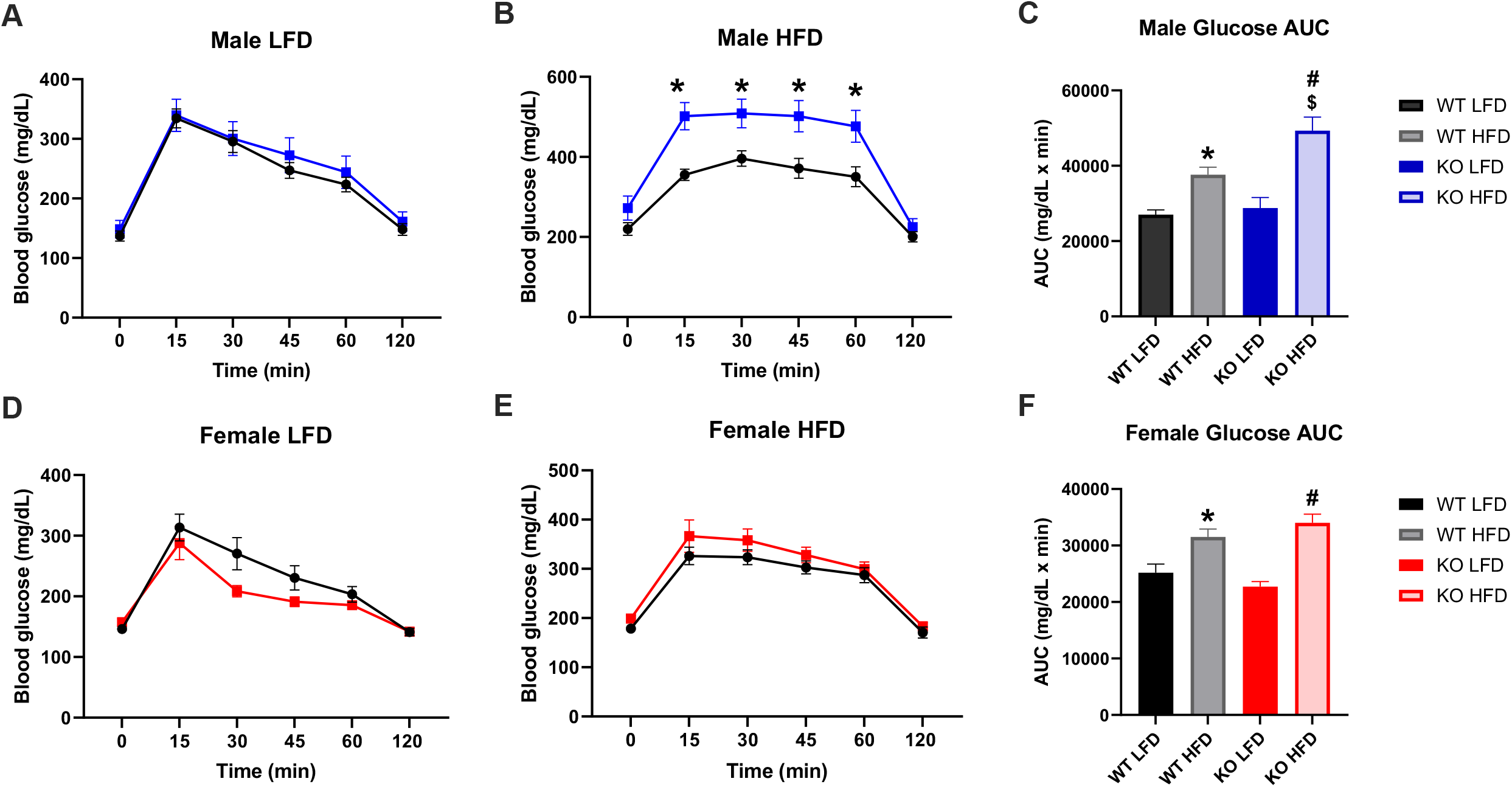
Glucose homeostasis is impaired in male GPR19 KO mice following exposure to a high fat diet. Male and female wildtype or GPR19 KO mice were fed a low-fat diet (LFD) and subject to an intraperitoneal glucose tolerance test (IPGTT). The same mice were then placed on a high-fat diet (HFD) for 10 weeks, after which a repeat IPGTT was performed. (A-C) LFD, HFD, and area under the curve (AUC) analysis for male mice. (D-F) LFD, HFD, and area under the curve (AUC) analysis for female mice. * = P < 0.05 vs WT LFD, # = P < 0.05 vs KO LFD, $ = P < 0.05 vs WT HFD, N = 7-9.

### 2.3 GPR19 deficiency results in increased liver mass in male mice after exposure to a high fat diet

More than 50 different GPCRs are predicted to be expressed in the mouse liver, of which most have been implicated in the regulation of liver metabolism ^[13] [19]^. Among these GPCRs includes the orphan receptor GPR19, however its specific role in hepatic physiology has yet to be explored. To assess this, we investigated the effects of GPR19 deficiency after high-fat feeding. Lipid analyses showed that GPR19 deficiency induced no significant changes in plasma triglyceride levels, however there was a non-significant trend towards a moderate decrease in liver triglyceride levels in female mice only (Fig. 3A,B,F,G). Male GPR19 KO mice exhibited a significant increase in absolute liver weight after a HFD (Fig. 3C), coupled with a moderate increase in liver weight to body weight ratio (Fig. 3D).

**Figure 3.**
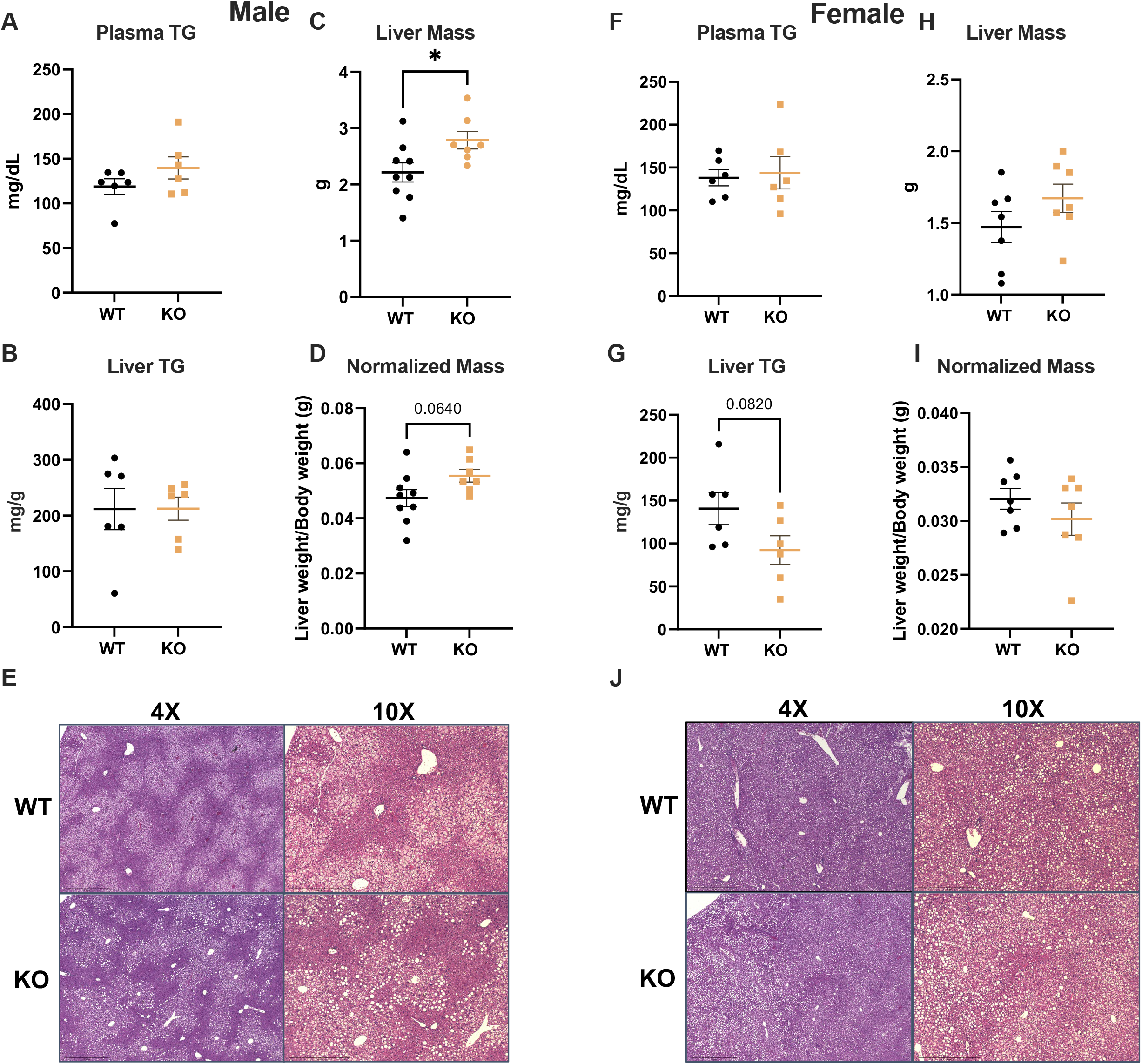
GPR19 deficiency results in increased liver mass in male mice after exposure to a high fat diet. After a 10 week HFD, male and female wildtype or GPR19 KO mice were analyzed for the following parameters: (A,F) plasma triglycerides, (B,G) liver triglycerides, (C,H) absolute liver mass, (D, I) liver mass normalized to total body weight, and (E,J) histology by H&E stain. * = P < 0.05, N = 6.

In contrast, no significant changes were seen in female liver weights compared to WT controls (Fig. 3H,I). Histological analyses by H&E staining showed no differences in hepatocellular injury in either male (Fig. 3E) or female mice (Fig. 3J) upon the loss of GPR19 expression. From these data we conclude that whole-body GPR19 deletion leads to a moderate, but significant sex-dependent hepatomegaly in mice after exposure to a high fat diet.

### 2.4 Whole-body GPR19 deficiency is associated with expression changes in hepatic lipid and glucose metabolism enzymes

We next investigated whether the increased liver weight observed in male GPR19 KO mice were linked to changes in hepatic metabolic enzymes. GPR19 KO mice fed a HFD displayed significant increases in *Pgc1a* (*Ppargc1a)* gene expression (a key regulator of energy metabolism) in males (Fig. 4A), with a trend towards increased expression in females (Fig. 4K), compared to WT controls. Paradoxically, there were significant decreases in fatty acid oxidation (*Cpt1a, Hadha*), triglyceride synthesis (*Dgat2*), gluconeogenesis (*Pck1*), and antioxidant (*Sod2*) gene expression in male GPR19 KO mice compared to WT controls (Fig. 4C,E,H-J). In female GPR19 KO mice, there were moderate decreases in *Pdk4* transcripts, with no differences in other glucose or lipid metabolism genes compared to WT controls (Fig. 4K-T).

**Figure 4.**
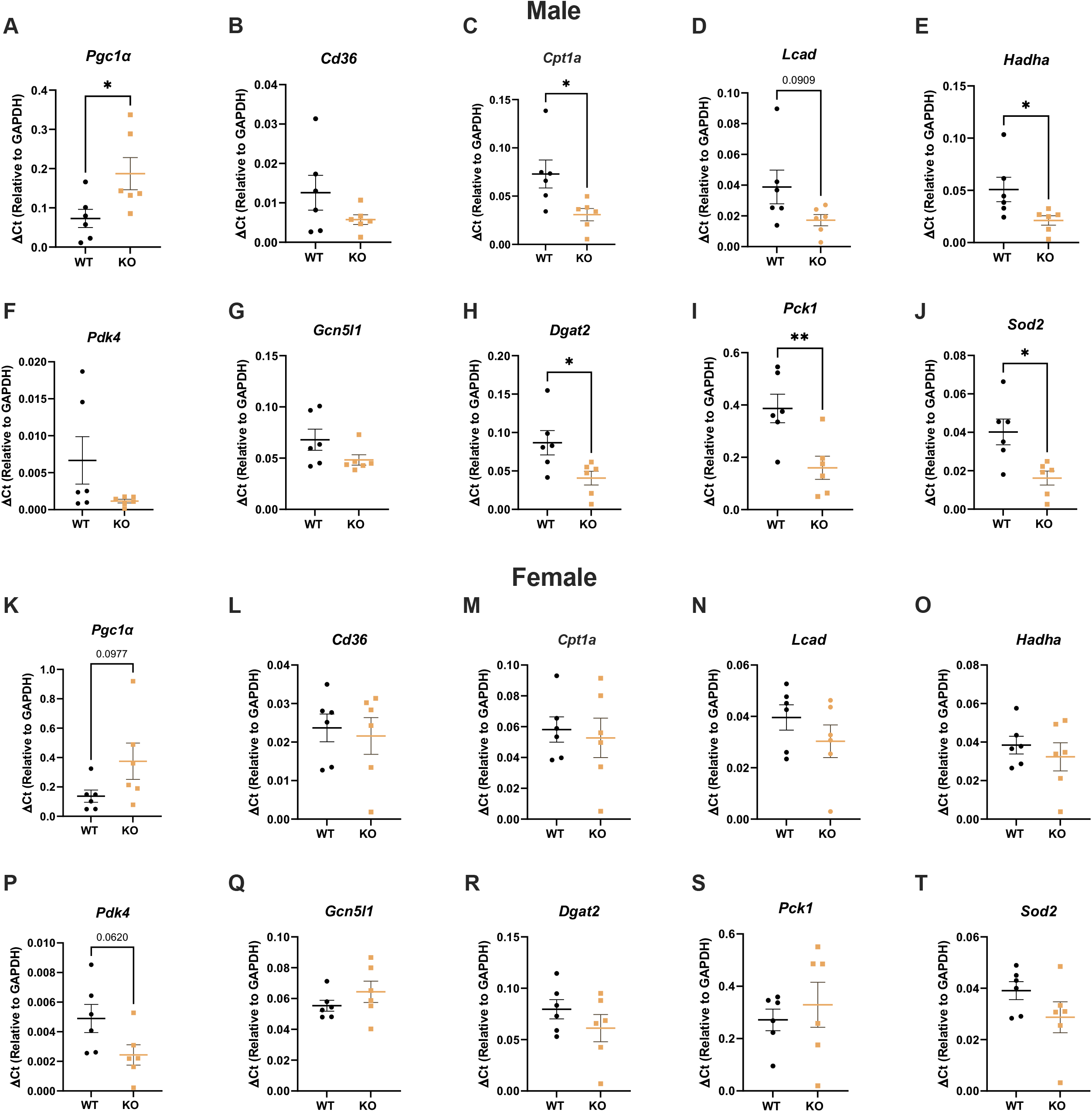
Whole-body GPR19 deficiency results in transcriptional changes in lipid and glucose metabolism genes. After a 10-week HFD, male (A-J) and female (K-T) wildtype or GPR19 KO mice were analyzed for the expression of hepatic metabolism genes. * = P < 0.05, ** = P < 0.01, N = 6.

We then investigated the protein abundance of key enzymes involved in hepatic metabolism by immunoblotting, and found significant decreases in the fatty acid uptake protein CD36, the acetylation of HADHA (Ac-HADHA; an inhibitory post-translational modification of hepatic FAO enzymes^; [20] [21]^, the hepatic gluconeogenesis enzyme glucose metabolism PEPCK, and the acetylation of the antioxidant enzyme, SOD2 (Ac-SOD2 - an inhibitory post-translational modification; ^[22]^) in male GPR19 KO mice compared to WT controls (Fig. 5A-M). In contrast, female GPR19 KO mice displayed significant decreases in fatty acid oxidation regulatory proteins (PGC1a, LCAD) and the glucose oxidation regulator, GCN5L1, with no differences in other fatty acid oxidation (CPT1a, CD36, Ac-HADHA, HADHA), glucose metabolism (PEPCK, G6PC, PDK4), or redox (SOD2) enzymes compared to WT mice (Fig. 5N-Z).

**Figure 5.**
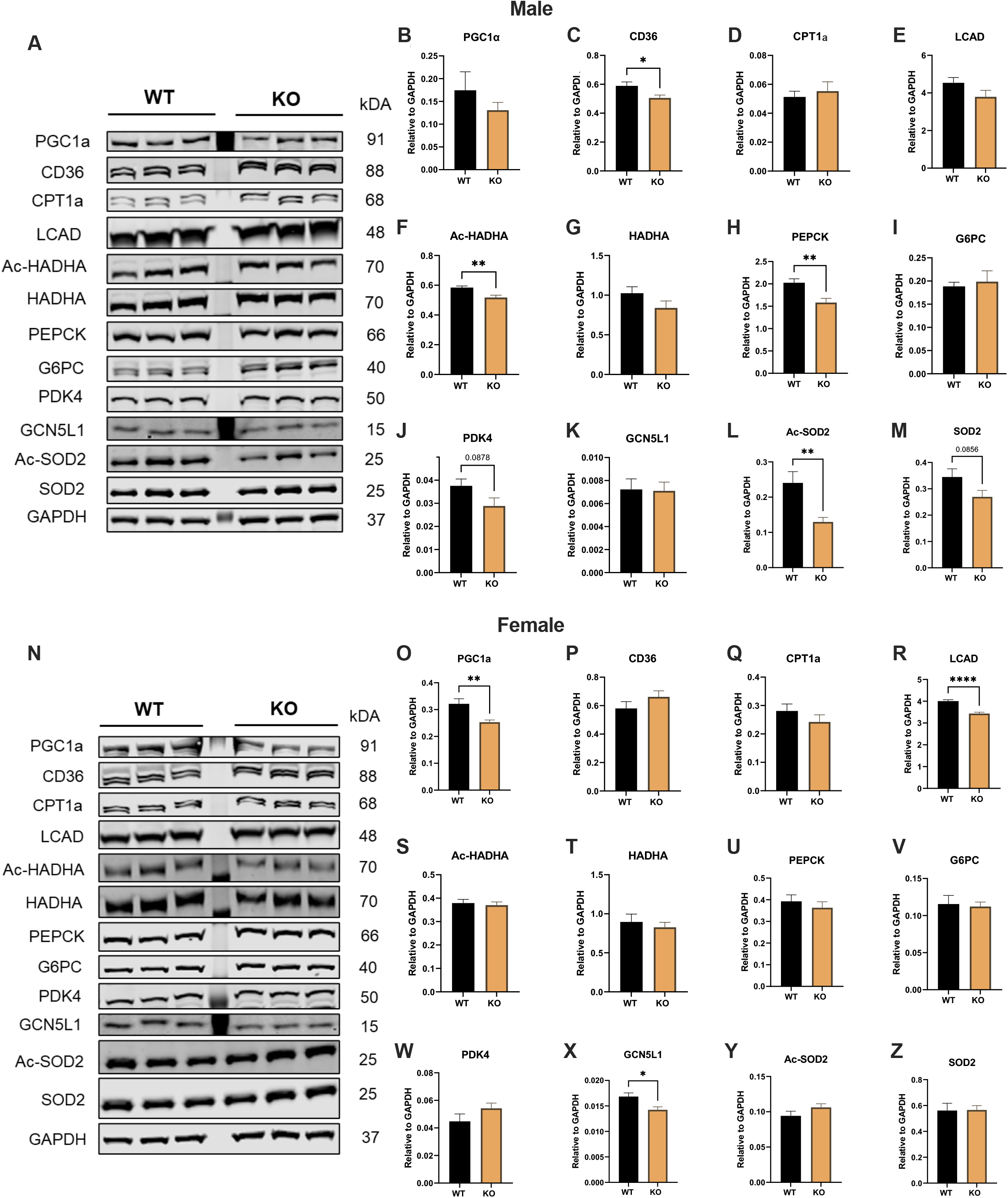
Loss of GPR19 is associated with changes in the protein abundance of lipid and glucose metabolic enzymes. After a 10-week HFD, male (A-M) and female (N-Z) wildtype or GPR19 KO mice were analyzed for the abundance of hepatic metabolism enzymes. * = P < 0.05, ** = P < 0.01, N = 6.

Taken together, these data suggest that whole-body GPR19 deficiency may disrupt hepatic metabolic homeostasis by decreasing *de novo* gluconeogenesis and reducing hepatic fatty acid oxidation in a sex-dependent manner.

## 3. DISCUSSION

In this study, we demonstrate for the first time a role for the orphan G-protein coupled receptor, GPR19, in the regulation of systemic metabolic homeostasis in diet-induced obesity. Using a novel whole-body *Gpr19*^*-/-*^ mouse model, we show that GPR19 deficiency induces a shift in whole-body energy metabolism by increasing energy expenditure and decreasing glucose disposal in a sex-dependent manner. Assessment of systemic metabolic performance suggests that GPR19 deficiency is associated with increased fat mass due to enhanced food intake and decreased physical activity. Male GPR19 KO mice show decreased whole-body glucose tolerance, combined with decreased expression of key hepatic glucose production enzymes, under obese conditions. Together, these findings reveal a potential role for GPR19 in mediating nutrient signaling and substrate oxidation preferences in the liver.

Since its initial discovery in the brain ^[10] [11]^, subsequent studies identified GPR19 as a circadian oscillating GPCR that alters locomotor activity rhythm ^[12]^ and regulates dipsogenic behavior in rodents ^[18]^. Our current data complements this behavioral phenotype in that GPR19 KO displayed a modest reduction in physical activity under both low-fat and high-fat fed conditions (Fig. 1I,J), suggesting a role in neural regulation ^[23] [24]^. Furthermore, the loss of GPR19 expression significantly increased food intake in male mice (Fig. 1G), which may suggest a potential link between thirst and hunger. The interdependency of these two behaviors is a long-established phenomenon, where one induces a concomitant response in the other ^[25] [26] [27]^. However, there are controversies on whether feeding or drinking is secondarily regulated ^[28] [29]^. Combined, these studies indicate that GPR19 may function in the regulation of energy homeostasis, however further work is required to establish the mechanism underlying its effects on feeding and drinking behavior.

Sex is fundamental factor in the maintenance of whole-body energy homeostasis ^[30]^. Although females are known to have a higher relative fat mass than males, they tend to be more glucose tolerant due to improved peripheral insulin sensitivity ^[31] [32] [33]^. In different rodent models of diet-induced obesity, males show a stronger phenotype than females, where the incidence of metabolic abnormalities is higher in males ^[34] [35] [36]^. This may explain the more pronounced effects of GPR19 deficiency on whole-body glucose disposal (Fig. 2) and tissue-specific metabolic flexibility (Fig. 4-5) observed in male mice compared to female mice. Stahlberg *et al*. ^[37]^ identified sexual dimorphic expression patterns in rodent livers, where males exhibited a higher expression of genes involved in hepatic fuel metabolism compared to females, indicating a greater capacity for regulating substrate use. This suggests that the males may be more sensitive to GPR19-dependent changes in the liver due to increased metabolic turnover. Further studies need to be conducted to investigate whether the effects of GPR19 on systemic and tissue-specific metabolism is due to intrinsic sexual dimorphism or other sex-specific features.

One caveat to our studies is that the animal model used, a whole-body GPR19 KO mouse, precludes the ability to rigorously interrogate the effects of GPR19 depletion in a tissue-dependent manner. As such, further work will be required to determine whether hepatocyte-specific loss of GPR19 drives the changes observed in lipid and glucose metabolism enzymes, or whether the decreased expression of triglyceride synthesis (*Dgat2*), fatty acid oxidation (*Cpt1a, Hadha*), and gluconeogenesis (*Pck1*) enzymes (Fig. 4) in HFD-fed male GPR19 KO mouse livers is a response to systemic changes in energy substrate metabolism. For example, decreased whole-body glucose tolerance may lead to a liver-specific downregulation of hepatic gluconeogenic enzymes, combined with a decrease in hepatic fatty acid oxidation (which provides the energy required to drive gluconeogenesis), as a means to limit hyperglycemia in HFD male GPR19 KO mice. This potential decrease in hepatic fatty acid oxidation may also lead to the increase in liver weight observed in these mice (Fig. 3), as it would impair the ability of the liver to reduce steatosis by upregulating fat use. Future work in isolated hepatocytes from GPR19 KO mice, or the generation of liver-specific GPR19 KO animals, should resolve this question.

A second caveat is that our studies did not address the role of adropin as a putative endogenous ligand for GPR19. Previous studies have shown that adropin, a liver- and brain-derived peptide hormone ^[38]^, acts as a ligand for GPR19 in several cell types ^[18] [39]^. Adropin levels are decreased in obesity, and restoration of normal adropin levels improves whole-body glucose homeostasis (see, e.g., ^[38] [40] [41]^. Future studies using our GPR19 KO mouse model will help to elucidate whether these effects are dependent on the actions of GPR19 *in vivo*.

In summary, our study provides robust evidence that GPR19 is a potential regulator of metabolic homeostasis in diet-induced obesity, and thus highlights this GPCR as a possible target for therapeutic applications in the progression of metabolic-associated diseases.

## 4. MATERIALS AND METHODS

### 4.1 Animal Care and Use

*Gpr19* heterozygous *(Gpr19*^+/−^) mice were obtained from the Mutant Mouse Resource & Research Centers (MMRRC strain name, C57BL/6N-*A*^*tm1Brd*^ Gpr19^*tm1b(KOMP)Mbp*^/JMmucd) (RRID:MMRRC_047948-UCD). Whole-body *Gpr19* null (*Gpr19*^−/−;^ KO) mice and wildtype (WT) controls were bred from heterozygote mutant crosses. Animals were housed in the University of Pittsburgh animal facility under standard conditions with *ad libitum* access to water and food and maintained on a constant 12-hour light/12-hour dark cycle. Male and female GPR19 WT and KO animals were fed a control low fat diet (LFD; 70% carbohydrate, 20% protein, 10% fat; Research Diets D12450B) from birth to 16-20 weeks of age, then switched to a high fat diet (HFD; 20% carbohydrate, 20% protein, 60% fat; Research Diets D12492), for 10 weeks. At the end of the 10 weeks, animals were euthanized by CO_2_ asphyxiation/cervical dislocation, and tissue/serum samples obtained for analysis. Experiments were conducted in compliance with National Institutes of Health guidelines and followed procedures approved by the University of Pittsburgh Institutional Animal Care and Use Committee.

### 4.2 Whole Body Composition and Metabolic Cage Studies

EchoMRI™ Body Composition Analysis System was used to measure whole body fat and lean mass in live animals. The Sable Systems Promethion Multiplexed Metabolic Cage System was used to evaluate physical activity, feeding, energy expenditure, and respiratory exchange ratio (RER) by indirect calorimetry. Studies consisted of 72-hour cage time, where the first 24 hours was considered acclimation, and the subsequent 48 hours was used to calculate the 24-hour and 12-hour light and dark cycle averages or totals for each parameter measured. LFD mice were analyzed immediately before the diet switch, and HFD-fed mice were studied in metabolic cages after 10 weeks of HFD feeding. Mice were individually housed during metabolic cage studies in compliance with IACUC-approved protocols.

### 4.3 Intraperitoneal Glucose Tolerance Test (IPGTT)

Glucose tolerance tests (GTTs) were performed as previously described ^[42]^ with minor modifications. After a 6-hour morning fast, basal plasma samples (*t* = 0) were collected, then mice were injected intraperitoneally with a 2 g/kg glucose bolus. Blood glucose was measured by tail bleed at set time points using a Bayer Contour Next EZ handheld glucometer (*t* = 15, 30, 45, 60, and 120 min).

### 4.4 Histology

Preparation and staining of all histological samples were conducted by the Pitt Biospecimen Core at the University of Pittsburgh. The right lobe of the liver was excised from each mouse following euthanasia, fixed in 10% buffered formalin phosphate overnight at shaking incubation, washed 3 times for 5 min with 1X phosphate-buffered saline (PBS), and then transferred to 70% ethanol. Samples were then embedded in paraffin for sectioning and stained with hematoxylin and eosin (H&E) for analysis. Histology sections were photographed using the Evos FL Auto 2 Microscope and representative images are shown.

### 4.5 Protein Isolation and Immunoblotting

Liver tissues were rapidly harvested following euthanasia, weighed and flash-frozen in liquid nitrogen. For protein isolation, tissues were minced and lysed in 1× CHAPS buffer (1% CHAPS, 150 mM NaCl, 10 mM HEPES, pH 7.4) using a VWR 4-Place Mini Bead Mill, then incubated on ice for ∼2.5 hours. Homogenates were spun at 10,000 *g* at 4°C for 10 min., and the supernatants collected for immunoblotting. For immunoblotting, protein lysates were quantitated using a BioDrop μLITE Analyzer, prepared in LDS sample buffer, separated using Bolt SDS-PAGE 4–12% or 12% Bis-Tris Plus gels, and transferred to nitrocellulose membranes (all Invitrogen). Membranes were blocked using Intercept (PBS) Blocking Buffer and incubated overnight in the following primary antibodies: rabbit PDK4 (PA5-13776) from ThermoFisher; goat PGC-1*α* (ab106814), rabbit acetyl K122-SOD2 (ab214675) from Abcam; rabbit GAPDH (2118S), rabbit SOD2 (D3×8F) from Cell Signaling Technologies; rabbit CD36 (18836-1-AP), rabbit CPT1a (15184-1-AP), rabbit CPT1b (22170-1-AP), rabbit HADHA (10758-1-AP), rabbit LCAD (17526-1-AP), rabbit PEPCK1 (16754-1-AP), and rabbit G6PC (22169-1-AP) from ProteinTech; HADHA acetyl-lysine (Lys-350/Lys-383/Lys-406) as reported previously ^[43]^; and GCN5L1 as previously reported ^[44]^. Protein loading was confirmed using GAPDH as a loading control. Fluorescent anti-goat or anti-rabbit secondary antibodies (red, 700 nm; green, 800 nm) from LiCor were used to detect expression levels. Images were obtained using Licor Odyssey CLx System and protein densitometry was measured using the LiCor Image Studio Lite Ver. 5.2 Software.

### 4.6 Lipid Analysis

Plasma and liver triglycerides (TG) were measured using the colorimetric Infinity Triglyceride Reagent Kit (ThermoFisher) and analyzed with the SkanIt™ Software (Ver. 6.0.2). Hepatic triglycerides were extracted using a methanol/chloroform-based method as previously described ^[45]^.

### 4.7 RNA Isolation and Quantitative RT-PCR

Total RNA was isolated from liver tissue using the RNeasy Mini Kit (Qiagen). RNA was quantitated, and 300 ng was used to generate cDNA using Maxima Reverse Transcriptase Kit (ThermoFisher). Quantitative real-time PCR was performed using 1× Power SYBR-Green PCR Master Mix (ThermoFisher), with validated gene-specific primers (Qiagen) on the QuantStudio™ 5 System. Each experiment was performed in triplicate and data was analyzed via the ΔC_t_ method using *Gapdh* as the reference gene.

### 4.8 Statistics

Means ± SEM were calculated for all data sets. Data were analyzed using two-way ANOVA with Tukey’s post-hoc testing to determine differences between genotypes and feeding groups. Time course data was analyzed using two-way ANOVA with Sidak’s post-hoc testing. Comparisons between single variable groups were made using unpaired, two-tailed Student’s T-Tests. *P* ≤ 0.05 was considered statistically significant. Statistical analyses were performed using GraphPad Prism 9.0 Software.

## 5. ACKNOWLEDGEMENTS

This work was supported by: National Institute of Health T32 Fellowship (T32GM133332) to B.A.S.M; and National Institute of Health research grant (R01HL147861) to I.S. The University of Pittsburgh Center for Metabolism and Mitochondrial Medicine is supported by the Pittsburgh Foundation (MR2020 109502) grant to M.J.J. This project used the University of Pittsburgh Medical Center (UPMC) Hillman Cancer Center and Tissue and Research Pathology/Pitt Biospecimen Core shared resource, which is supported in part by award P30CA047904.

## 6. AUTHOR CONTRIBUTIONS

M.J.J. and I.S. conceived the study and designed experiments; B.A.S.M, B.X., I.J.S., A.M.V., and M.W.S. performed experiments; B.A.S.M., B.X., M.J.J., and I.S. analyzed data; B.A.S.M., M.J.J., and I.S. prepared figures; D.T., J.R.M., and P.B. provided critical input and expertise; B.A.S.M. drafted manuscript; B.A.S.M., M.J.J., and I.S. edited and revised manuscript.

